# Unifying transcranial focused ultrasound and transcranial magnetic stimulation effects with calcium-dependent synaptic plasticity theory

**DOI:** 10.64898/2026.07.09.737614

**Authors:** Yupeng Tian, Kevin Kadak, Khushi Kankaria, Ravindu Upasena, V. Kumar Murty, Robert Chen, John D. Griffiths

## Abstract

Low-intensity transcranial focused ultrasound stimulation (TUS) is an emerging technology that shares features of both established invasive neurostimulation techniques such as deep brain stimulation (DBS) and noninvasive techniques such as transcranial magnetic stimulation (TMS). Like DBS, TUS can target non-superficial brain structures with millimeter-level precision. Like TMS, but unlike DBS, the most important physiological effect of TUS from a clinical perspective is its ability to induce lasting neuroplastic changes (long term potentiation/depression; LTP/LTD) from relatively short stimulation sessions. Thus follows the intriguing possibility that, although TUS and TMS have fundamentally different primary mechanisms of action – acoustic versus electromagnetic – they might nevertheless share a common secondary mechanism of plasticity induction through temporally patterned stimulation. A quantitative mathematical theory of this secondary mechanistic pathway could therefore have important explanatory and predictive value in both modalities. Two major challenges to the development of such a theory, however, are: i) experimental results showing contradictory plasticity effects between TUS and TMS for nominally similar stimulation parameters, and ii) the markedly different temporal structures of their stimulation waveforms (ranging from discrete pulses in TMS to continuous sinusoidal oscillations in TUS), even for highly aligned protocol designs such as continuous theta burst (cTB) stimulation. Here we show that a mathematical model of calcium-dependent synaptic plasticity in corticothalamic circuits, already developed extensively for TMS, can indeed provide such a unified description of stimulation effects across these two modalities. Numerical simulations using this model for a range of TUS and TMS protocols reproduced plasticity effects consistent with experimentally observed changes in cortical excitability. In particular, our model addresses both of the above challenges, by i) reconciling apparently contradictory results across modalities for the same stimulation parameters, and ii) introducing a simple algebraic approach, which we term the ‘equivalent energy principle’, for relating corresponding TMS and TUS waveforms. The ability of the model to account for differing effects across multiple stimulation modalities provides further support for the underlying general theory describing calcium-based regulation of stimulation plasticity effects, which spans multiple scales of system organization – from ion channel kinetics to neural population activity. Our work also provides a foundation for future bidirectional exchange of new experimental observations and insights between experimental and theoretical TUS and TMS research, including strategies for model-based protocol optimization and discovery of novel plasticity-inducing TUS and TMS paradigms.

## Introduction

Transcranial focused ultrasound stimulation (TUS) is a relatively new clinical neurotechnology that in recent years has become a major focal point of research and development in the field of brain stimulation. Both the high-intensity (ablative [1]) and low-intensity (sonoporational [2] and neuromodulatory [3]) variants of TUS have shown great promise as non-surgical and non-pharmacological therapeutic options for treatment-resistant patients presenting with a range of psychiatric and neurological disorders, including Parkinson’s disease [3], essential tremor [1], brain tumor [4], Alzheimer’s disease [2], depression [5], and obsessive compulsive disorder [6]. The main physical effects of TUS on biological structures include thermal effects [7], cavitation (gas bubble formation) [2], radiation wave propagation [8] and mechanical effects on biological membrane [9]. Scientific understanding of how exactly these primary mechanisms of action impact neural systems is still at a nascent stage however, and may differ substantially across the three main TUS variants.

For neuromodulatory applications, experimental and theoretical modeling work has explored multiple candidate pathways, including local pressure wave propagation [10], membrane potential change [7], intramembrane cavitation [11], and heating [12]. In particular, the general physical and immediate neuron-level physiological effects of TUS have received intense interest in recent years, from both the computational and the broader neuroscientific community. However the (arguably more important) question of how exactly TUS modulates *plasticity* (synaptic or otherwise), and in turn induces measurable long-term effects on brain activity, to date remains almost entirely unexplored.

Transcranial magnetic stimulation (TMS) is an established noninvasive therapeutic technique that is an effective second-line treatment for depression [13, 14], obsessive compulsive disorder [15], addiction [16], and stroke [17], and under active investigation for a wide range of other conditions. TMS works by activating neurons via secondary electrical currents generated in the brain, perpendicularly to the transcranially-applied primary magnetic field from a scalp-positioned coil [18]. Unlike TUS, TMS cannot reach deep brain structures, and has inherently lower spatial resolution [19, 20]. Nevertheless, a critical question for the field remains whether stimulation targeted at those superficial cortical regions that are accessible by both modalities has similar physiological effects, particularly when using comparable temporal stimulation protocols and intensity levels. Such a correspondence, if discovered, could be extremely valuable - allowing the extensive experience accumulated from over 30 years of clinical and basic human neurophysiology research using TMS to be drawn on directly by TUS researchers and medical practitioners. One of the key insights from this three-decade period is the observation that different TMS stimulation patterns have different plasticity effects. For example, with regular repetitive TMS (rTMS), in which stimulation consists of repetitive pulses without burst structures, low-frequency (∼1Hz) stimulation generally reduces cortical excitability and is widely considered to induce LTD-like plasticity [21], whereas higher frequencies (10+ Hz) typically increase cortical excitability and are associated with LTP-like plasticity [22].

The most extensively studied and utilized rTMS protocol class in the ∼20 years since its invention is continuous/intermittent theta-burst (c/iTB) stimulation. TB delivers stimulation in structured bursts with specified intra- and inter-burst frequencies (typically 30-50Hz and 5Hz, respectively), and a fixed number of pulses-per-burst (typically 3-5). iTBS adds to this a slow ON/OFF cycle, typically comprising 2*s* ON periods, alternating with 8*s* OFF periods. The iTB-TMS-200s protocol, which delivers approximately 600 pulses over 200*s*, is a typical facilitatory protocol that induces LTP-like plasticity. It has been applied clinically for the treatment of major depressive disorder [23], post-stroke rehabilitation [17], and Parkinson’s disease [24]. Unlike iTB, cTB applies bursting stimulation continuously, and therefore requires far shorter session durations to deliver the same number of pulses. The most commonly used cTB protocol, cTB-TMS-40s, which delivers the same number of pulses as iTB-TMS-200s but over a much shorter session duration, is a well-established inhibitory protocol that generally induces LTD-like plasticity [25]. It has been investigated in clinical trials such as for the treatment of obsessive-compulsive disorder [15, 26]. Interestingly, extending the stimulation duration to 80*s* (cTB-TMS-80s), but keeping all other parameters identical, has been reported to induce an LTP-like effect in MEPs – inverting the sign of the textbook plasticity response for the more common cTB-TMS-40s [25, 27].

cTB has also recently been developed for TUS applications [3, 28]. In cTB-TUS protocols, the pulse/burst repetition frequency is set to 5*Hz*, with stimulation delivered continuously without intervals, corresponding to complete TUS-OFF periods [28, 29]. cTB-TUS has been shown to increase cortical excitability in the left primary motor cortex (M1) when stimulation is applied to left M1 [28]. In addition, cTB-TUS has been applied to the globus pallidus internus (GPi) within the basal ganglia to investigate therapeutic effects in patients with Parkinson’s disease and dystonia [3]. This study demonstrated that cTB-TUS significantly increases theta-band power in GPi local field potentials, and that the increase in theta power is consistent with the therapeutic effects observed with dopaminergic medications in Parkinson’s disease [3]. Although these protocol-specific plasticity effects are widely reproducible, the response to theta-burst stimulation may vary across individuals and experimental conditions for both TMS and TUS, with the same protocol occasionally producing reduced, absent, or opposite effects from the canonical LTP- or LTD-like response [30, 31].

Despite this large amount of experimental data and paradigms across TMS, TUS, and related neurostimulation modalities, the precise mechanisms underlying stimulation-induced plasticity responses remains an active area of research in pre-clinical, human neurophysiology, and computational neuroscience research [32–35]. For several decades it has been known that intracellular calcium concentration is a critical factor governing neuronal plasticity induced by external stimulation [36, 37]. Specifically, synaptic plasticity is regulated by intracellular calcium levels relative to depression and potentiation thresholds: long-term potentiation (LTP) is induced when intracellular calcium concentration exceeds the latter, whereas long-term depression (LTD) is induced when calcium concentration lies between the two [38, 39]. This nonlinear relationship between intracellular calcium concentration and plasticity induction is often characterized using an inverted-Ω-shaped function [40, 34]. Based on this formulation, calcium-dependent plasticity (CaDP) theory has been incorporated into mathematical models for mechanistic investigations of plasticity induced by stimulation with direct current injection [39] and with TMS [33, 34, 41–43]. These models have proved remarkably effective at reproducing and explaining, with minimal ad-hoc parameter tuning, a wide range of both long-term and short-term experimental plasticity effects – including rTMS, iTB-TMS, cTB-TMS, and paired-pulse short- and long-interval intracortical inhibition and facilitation [34, 41–43, 33, 44–46, 35, 47].

Motivated by these prior successes, and by the compelling possibility of a general mathematical theory for neurostimulation-induced plasticity, the present work proposes a minimal modification of the TMS modelling approach used previously by ourselves and others, extending it naturally into the domain of TUS. Our aims were twofold: i) to successfully replicate (again, with minimal ad hoc parameter tuning) published experimental TUS plasticity results, and ii) to provide new insight into as-yet unresolved questions on the relationship between TMS and TUS findings, which show several notable inconsistencies. Following the approach of Kadak et al. (2025a,b) [33, 44], Wilson et al. (2018, 2021) [48, 47], and Fung and Robinson (2013, 2014) [34, 41], we developed a minimal mean-field circuit model comprising cortical and thalamic neural populations, with physiologically realistic endogenous population-level oscillatory activity, CaDP of *α*-amino-3-hydroxy-5-methyl-4-isoxazolepropionic acid (AMPA) receptors mediated by time-varying N-methyl-D-aspartate (NMDA) receptor conductances, and Bienenstock-Cooper-Munroe (BCM) metaplasticity. We introduce a new approach for defining consistent amplitude scaling between modeled TMS and TUS waveforms (termed the *equivalent energy principle*; **Fig. 1**), allowing us to evaluate the unique effects of TMS vs. TUS temporal waveform structure.

**Figure 1:**
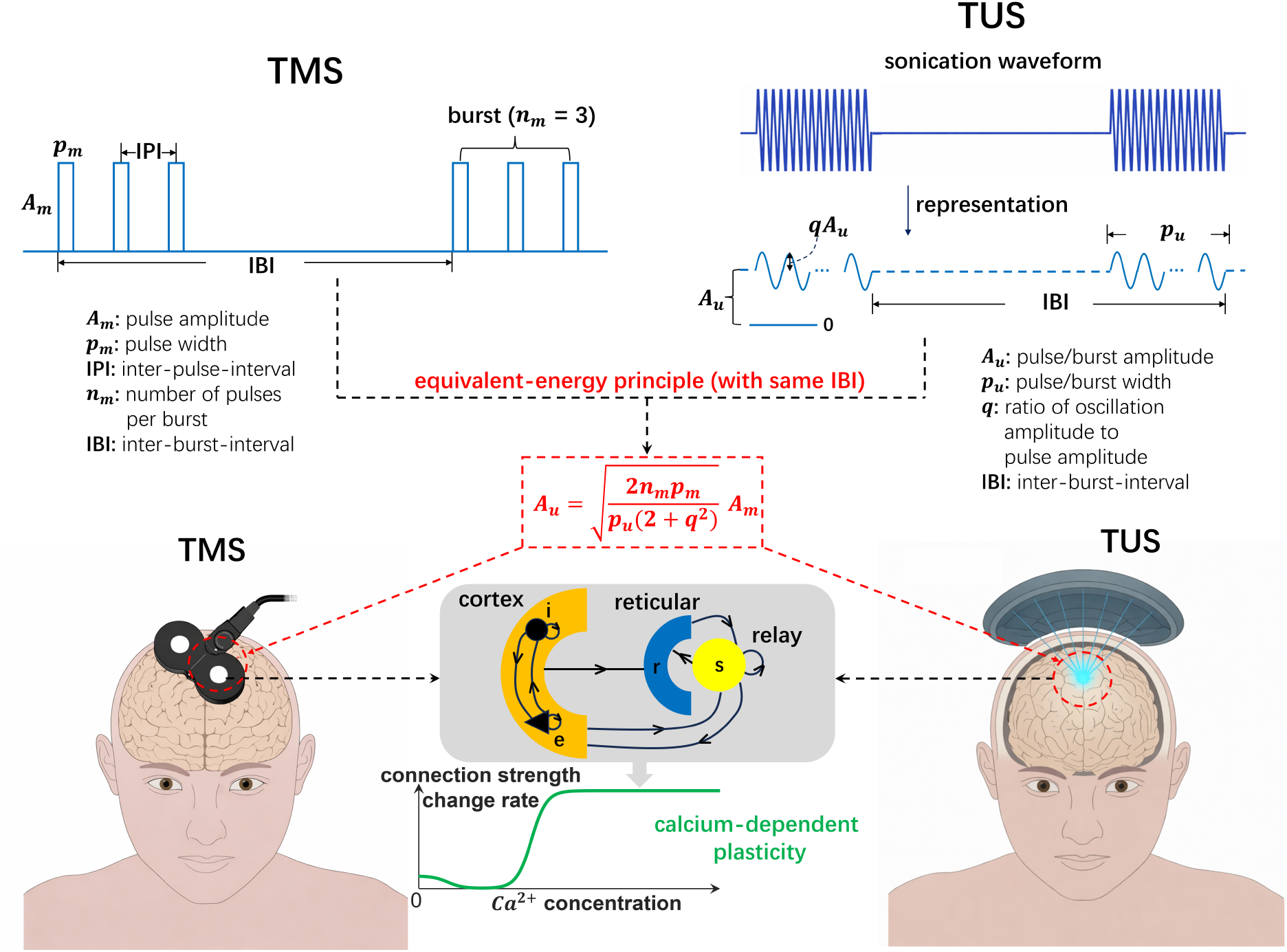
Unified TUS/TMS model schematic and equivalent-energy principle formulation and application. *Top:* Stimulation waveforms used in complex TMS protocols such as cTB consist of structured sequences of bursts of discrete pulses. In contrast, TUS protocol waveforms use smoothly varying bursts/pulses. In our model, the original *MHz*-frequency TUS waveform is represented as a sub-*KHz*-frequency sinusoid, placing its frequency structure within the range that neurons are responsive to. *Middle:* Parameters of TMS and TUS bursts are linked through the proposed equivalent-energy principle (red box; **Eq. 1**), which specifies the parameter relationship required for the two stimulation modalities to deliver the same amount of energy. Solving for *A*_*u*_ determines the implied TUS amplitude scaling factor with no free parameters. *Bottom:* The TUS/TMS waveform is then delivered as an afferent input to the excitatory neural population of a standard corticothalamic circuit, including cortical populations (excitatory and inhibitory neurons), thalamic populations (relay and reticular nuclei), and their synaptic connections. The calcium-dependent synaptic plasticity mechanism is active in the stimulated population only, and identical circuit parameters are used for both stimulation modalities (**Table S1**). Brain+device figures modified from [2]. See main text for abbreviation definitions.

Our principal explanatory target is published data from several experimental studies [29, 25, 27] showing pre vs. post M1 stimulation change in MEP amplitudes for different durations and frequencies of TMS and TUS stimulation, totaling 9 unique stimulation protocol-modality combinations, including cTB-TMS-40s, cTB-TMS-80s, cTB-TUS-40s, cTB-TUS-80s, iTB-TMS-200s, rTMS-1Hz-600s, rTMS-20Hz-30s, TUS-2Hz-80s and TUS-10Hz-80s. Simulated stimulation effects were qualitatively and quantitatively consistent with the majority of these reported values, including several notable sign-changes from LTD to LTP, arising from dosage-frequency interactions. This work provides a foundational framework and starting point for understanding TUS-induced plasticity mechanisms, and demonstrates strong potential for future optimization of TUS stimulation parameters to improve therapeutic outcomes.

## Results

### An equivalent-energy principle for joint parameterization and modeling of cTB-TUS/TMS waveforms

As a first step in building a unified cTB-TUS/TMS modeling framework, it was necessary to develop an approach for parameterizing cTB stimulation waveforms across the two modalities in a consistent way. To do this, we constructed an expression (**Eq. 1** and **Fig. 1**; see full description in **Methods** and full derivation in **Supplementary Methods**) that provides, for a given set of cTB-TUS and cTB-TMS waveform parameters, a scaling factor *A*_*u*_ for which the same amount of energy is delivered in both modalities. This operation, which we term the *equivalent energy principle* is an ansatz with which we begin our analysis, representing the hypothesis that differences in TMS and TUS plasticity effects can be explained by waveform shape alone (discrete rectangular pulses vs. sinusoidal waves). Whilst our parameterization is specific to cTB, we note that both the equation and the concept of maintaining equivalent energy between two protocols is quite general, and can also readily be used outside of the present mathematical modeling context.

### Calcium-dependent plasticity model accurately reproduces TMS and TUS dosage and frequency effects

Numerical simulations using our new corticothalamic circuit-based plasticity model (red bars) and the original Fung-Robinson model (blue bars), alongside reported experimental results (white bars) from 10 different TMS and TUS protocols, are shown in **Fig. 2**. In the computational models, plasticity was quantified as the percentage change in synaptic strength (*ν*_*ee*_) within the targeted population from pre-to post-stimulation (**Eqs. 18**-**19, Fig. 2a**). In the empirical data, plasticity was similarly quantified as the percentage change in MEP amplitudes from the pre-stimulation baseline [25]. **Figs. 2b and 2c** compare the plasticity effects predicted by our model with empirical cTB-TMS and cTB-TUS data reported by Gamboa et al. [25] and Zeng et al. [29] respectively, as both the original values (**Fig. 2c**) and after normalization to the strongest effect (**Fig. 2b**). The model results are in excellent agreement with empirical data, although better with normalized values for TMS and better with original values for TUS. The remaining amplitude mismatch between the model predictions and the empirical observations may arise because the modeled change in cortical excitability alone is insufficient to fully account for MEP modulation, which is additionally influenced by downstream corticospinal and muscular pathways [50, 51].

**Figure 2:**
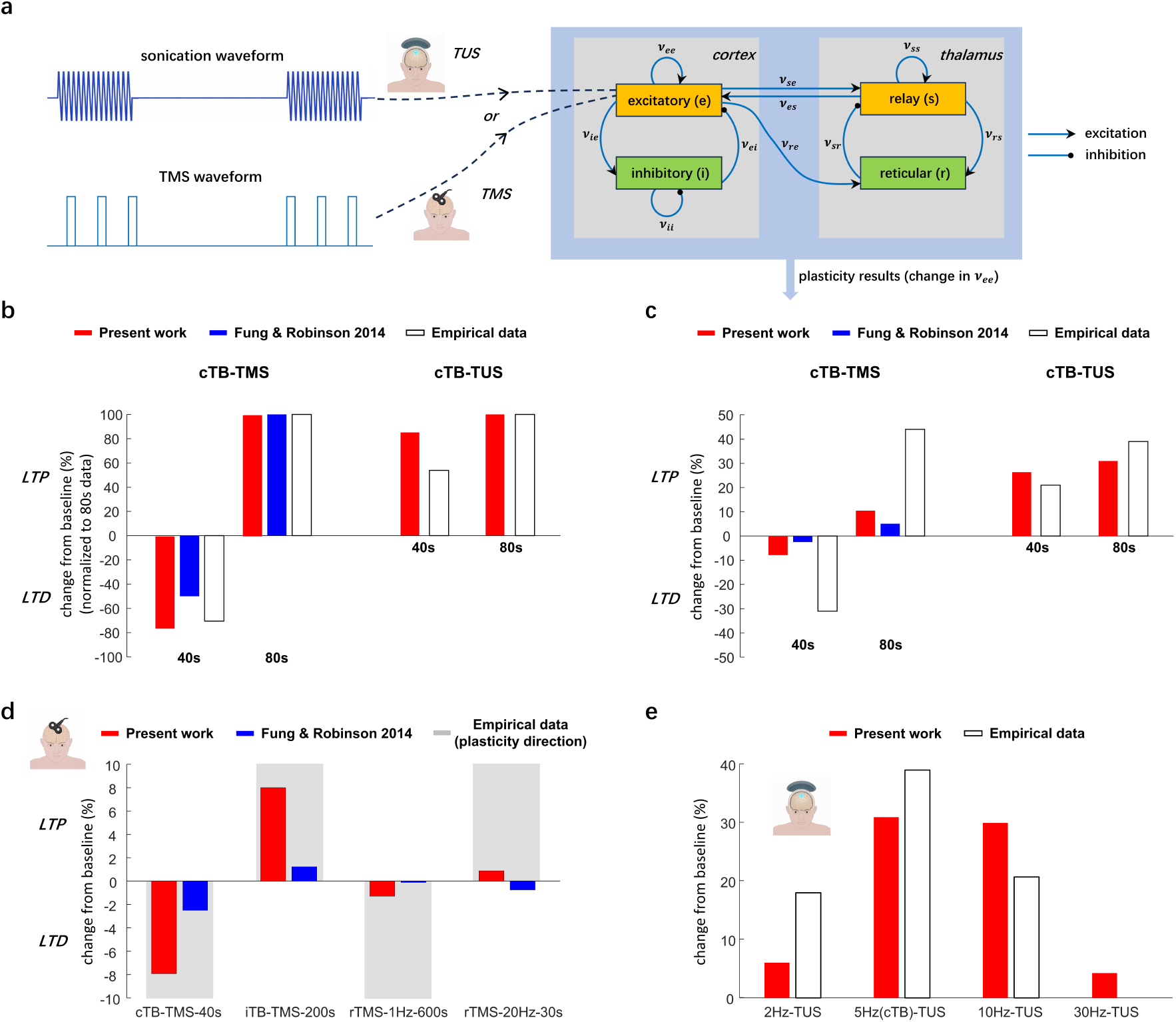
Unified TUS-TMS model captures multiple empirical stimulation-induced plasticity effects. Model predictions are compared with empirical data from TMS and TUS studies, as well as with the model of Fung and Robinson (2014) [41]. **a)** Schematic of the modeling framework and comparison of model-predicted and empirical plasticity for cTB-TMS and cTB-TUS protocols. TMS and TUS waveforms are applied to the corticothalamic model, and stimulation-induced plasticity is quantified by the change in the cortical excitatory synaptic strength (*ν*_*ee*_). **b-c)** Model predictions are compared with empirical MEP changes for cTB-TMS and cTB-TUS protocols with stimulation durations of 40 s and 80 s. TMS data are from Gamboa et al. (2010) [25] (MEPs recorded 10 min post-TMS), and TUS data are from Zeng et al. (2024) [29] (MEPs recorded 5 min post-TUS). **d)** Comparison under additional TMS protocols besides cTB-TMS-40s, including iTB-TMS-200s, rTMS-1Hz-600s, and rTMS-20Hz-30s. Stimulation durations were adjusted so that all protocols delivered the same number of pulses. Model predictions are compared with empirical results from Gamboa et al. (2010) [25], Chung et al. (2017) [49], Chen et al. (1997) [21], and Pascual-Leone et al. (1994) [22], respectively. **e)** Comparison under additional TUS protocols with PRFs of 2, 5, and 10 Hz [29]. Duty cycle (10%), sonication duration (80 s) and TUS amplitude (= 10.72, determined from the equivalent-energy mapping from the reference cTB protocol (**Eq. 1**)) were held constant. The 30 Hz result is a model prediction, as no corresponding experimental data are available.

There are two highly important properties of these data that are successfully captured by the model. First, the reversal in plasticity direction for cTB-TMS when protocol duration is doubled (LTD for 40s and LTP for 80s) is striking. A similar sign-flip effect has also been reported for iTB-TMS (both in data [25] and in simulations [41]) when the number of pulses is doubled from 600 to 1200. Second, instead of showing a sign-flip like the cTB-TMS, cTB-TUS shows a linear increase in plasticity magnitude with doubling protocol duration. Given that the total energy received in the two protocols is constrained by design to be identical, these results highlight the fact that even relatively subtle waveform differences (such as the difference between cTB-TUS and cTB-TMS) are sufficient to force dramatic shifts in plasticity responses.

Next, we further validated the model by comparing its predictions with experimental observations across additional stimulation protocols: iTB-TMS-200s [49], rTMS-1Hz [21], and rTMS-20Hz [22] (**Fig. 2d**). Again, a crucial property of these experimental data that is also accurately captured by our model are the sign flips, occurring as a function of carrier frequency (LTD for rTMS 1Hz vs. LTP for 20Hz) and intermittency (LTD for cTB-TMS vs. LTP for iTB-TMS). Additionally, our model correctly predicted that, for the same number of delivered pulses, TB-TMS protocols induce larger plasticity effects than rTMS protocols. This observation is also consistent with reports that TB-TMS elicits robust cortical plasticity and achieves clinical efficacy comparable to that of rTMS, while requiring substantially shorter treatment sessions [52, 53]. In both these rTMS vs. TB-TMS comparisons and in the cTB-TMS comparisons of **Fig. 2c**, the superior performance of our model over the Fung-Robinson model (on which it is closely based) is likely due to the inclusion of more detailed neuronal circuitry, including the cortical inhibitory population and thalamic nuclei, which are absent in the simpler one-population model of Fung et al. [41].

The final set of model validations are shown in **Fig. 2e**, where we simulated the additional TUS data reported by Zeng et al. [29], in which MEP responses were recorded under different pulse repetition frequencies (PRFs), in addition to the 5*Hz* condition used in cTB-TUS. Zeng et al. observed reduced LTP effects - represented by smaller MEP changes from pre-to post-stimulation - when the PRF deviated from 5*Hz* towards either 2*Hz* or 10*Hz* (**Fig. 2e**) [29]. Our model reproduces the experimentally observed PRF dependence, with the strongest plasticity response occurring around 5*Hz* and weaker responses at PRFs away from this condition (**Fig. 2e**).

### Intracellular calcium flux and depression/potentiation thresholds determine stimulation response patterns

The results of **Fig. 2** establish strong correspondence of our model with multiple empirical observations, across both stimulation modalities – including, most notably, accurate prediction of plasticity direction flips as a function of protocol. Accurately replicating empirical data *per se* is neither explanatory nor interesting, however; for this we must take a closer look at the internal workings and hypothesized mechanisms of the simulated system. **Fig. 3** shows a breakdown of the state variable time series during simulated TUS and TMS experiments, with a focus on the CaDP mechanism (**Fig. 3a** and **Fig. 3b, Eqs. 7-17**). In this framework, intracellular calcium concentration [*Ca*^2+^] regulates synaptic plasticity through the calcium-control Ω-function, producing LTD at moderate calcium levels and LTP at higher calcium levels (**Fig. 3c**). The resulting change in synaptic strength is represented by the stabilized connectivity *ν*(*t*), which is obtained from the instantaneous synaptic strength 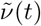 through a low-pass filtering process. Calcium dynamics are further modulated by BCM metaplasticity through NMDA conductance *g*_*NMDA*_(*t*) **Fig. 3b**, providing the mechanistic basis for the protocol-dependent plasticity shown in **Fig. 3e**.

**Figure 3:**
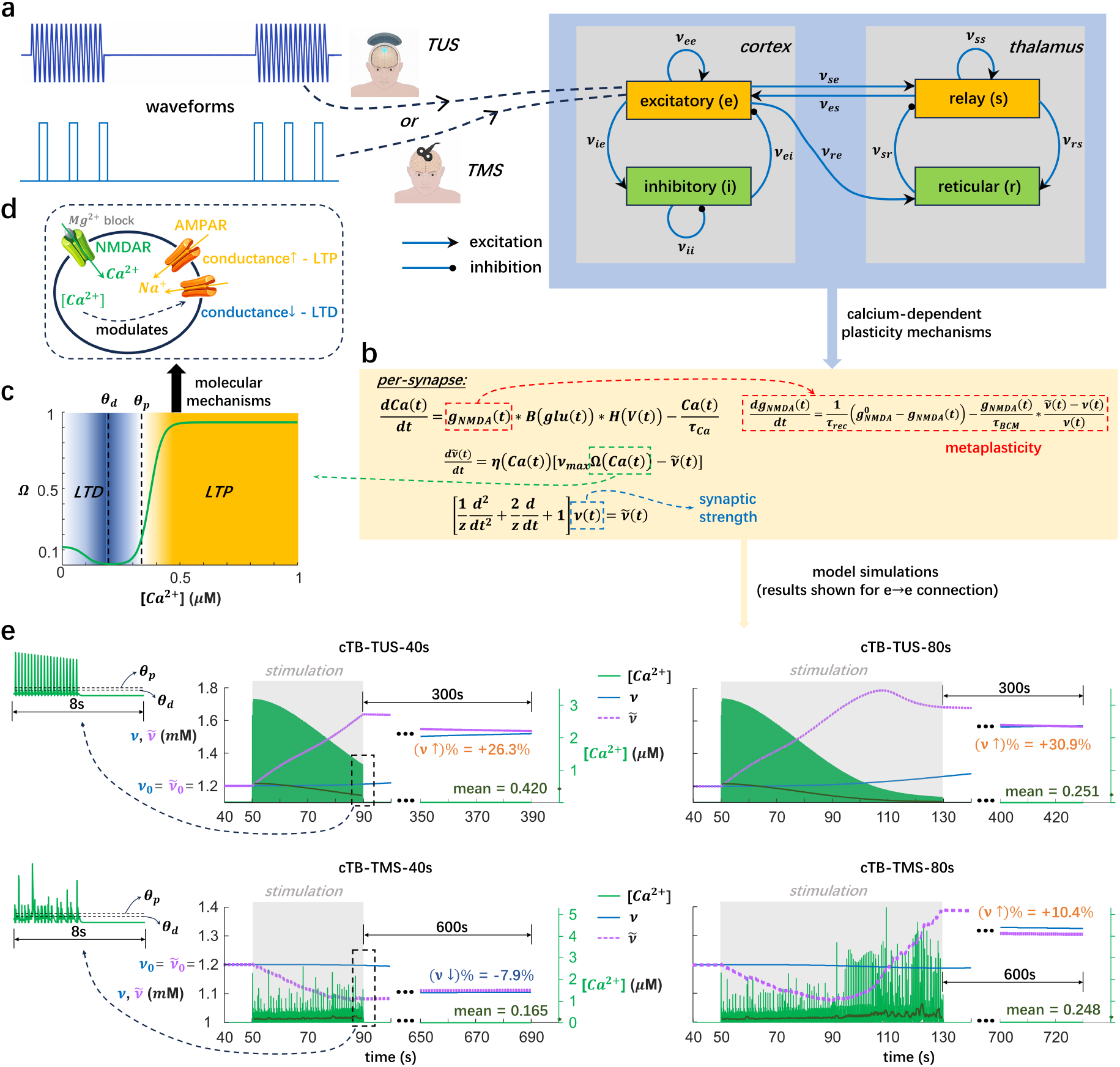
Mechanistic explanation of variable plasticity outcomes across stimulation protocols and modalities. **a)** Ultrasonic and electromagnetic stimulation waveforms are applied to a corticothalamic network comprising four neural populations: cortical excitatory (*e*) and inhibitory (*i*) neurons and thalamic relay (*s*) and reticular (*r*) nuclei. *ν*_*ab*_ denotes the synaptic connection strength from population *b* to population *a*. **b)** Model equations describing calcium-dependent plasticity (CaDP). **c)** The Ω-function relates intracellular calcium concentration ([*Ca*^2+^]) to synaptic modification, with depression (*θ*_*d*_) and potentiation (*θ*_*p*_) thresholds defining calcium ranges associated with LTD and LTP. **d)** Molecular interepretation of the CaDP mechanism: calcium influx is mediated primarily by NMDA receptors and modulated by extracellular *Mg*^2+^, while intracellular [*Ca*^2+^] regulates AMPA receptor conductance associated with the expression of synaptic plasticity. **e)** CaDP dynamics underlying the different plasticity outcomes of cTB-TUS-40s, cTB-TUS-80s, cTB-TMS-40s, and cTB-TMS-80s shown in **Fig. 2**. The more continuous TUS waveform produces sustained calcium elevations: during cTB-TUS-40s, [*Ca*^2+^] remains predominantly within the LTP-inducing range (mean [*Ca*^2+^] = 0.420*µM* ), driving an increase in the instantaneous synaptic modification 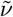 and ultimately in the stabilized synaptic strength *ν*. During cTB-TUS-80s, calcium initially occupies the LTP-inducing range before shifting towards the LTD-inducing range later in stimulation (mean [*Ca*^2+^] = 0.251*µM* ); nevertheless, the accumulated increase in 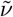 results in a net post-stimulation LTP effect. In contrast, the discrete TMS pulses produce intermittent calcium transients. During cTB-TMS-40s, calcium remains predominantly within the LTD-inducing range (mean [*Ca*^2+^] = 0.165*µM* ), decreasing 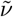 and ultimately *ν*. Extending cTB-TMS to 80*s* produces larger calcium elevations during the later phase of stimulation, shifting synaptic modification towards potentiation and ultimately reversing the post-stimulation outcome from LTD to LTP. Thus, the distinct temporal calcium dynamics induced by TUS and TMS provide a mechanistic explanation for their different duration-dependent plasticity outcomes. The dark-green traces show the 1*s* moving average of [*Ca*^2+^], and the overall mean calcium concentration during stimulation is indicated in each panel.

Plasticity is quantified by the percentage change in the *e* → *e* synaptic strength, *ν*_*ee*_. **Fig. 3e** shows representative simulations for four experimentally validated protocols: cTB-TUS-40s, cTB-TUS-80s, cTB-TMS-40s, and cTB-TMS-80s. Intracellular calcium dynamics regulate the evolution of the instantaneous synaptic strength 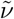 through the calcium-control functions *η* and Ω (**Eqs. 14-17**). The dark green trace shows the 1-s moving average of intracellular calcium concentration. During cTB-TUS-40s, the moving average of [*Ca*^2+^] remains within the LTP region of the Ω-function, with an overall mean [*Ca*^2+^] of 0.420*µM* (**Fig. 3e**). This behavior is consistent with the monotonic increase of 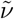 from its initial value 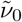 during cTB-TUS-40s. The increase in 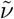 subsequently drives an increase in the stabilized synaptic strength *ν* from its baseline value *ν*_0_ through the low-pass filtering process (**Fig. 3b** and **Fig. 3e**). The difference between 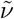 and *ν* additionally modulates [*Ca*^2+^] through the metaplasticity dynamics mediated by *g*_*NMDA*_ (**Fig. 3b** and **Fig. 3e**). During cTB-TUS-80s, the moving average of [*Ca*^2+^] remains within the LTP region during the initial ∼ 50*s* and subsequently shifts into the LTD region for the remainder of the stimulation period (**Fig. 3e**). Consequently, 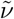 begins to decrease after reaching its peak value. Nevertheless, after the full 80*s* stimulation period, the net increase in 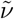 remains greater than that obtained with cTB-TUS-40s (**Fig. 3e**). As a result, after 300*s* post-TUS, the stabilized synaptic strength *ν* reaches a larger increase for cTB-TUS-80s (30.9% above baseline) than cTB-TUS-40s (26.3% above baseline) (**Fig. 3e**), indicating a stronger LTP effect, consistent with the empirical observations reported by Zeng et al. [29]. A reversal in plasticity direction is observed when comparing the simulations of cTB-TMS-40s (LTD) and cTB-TMS-80s (LTP) (**Fig. 3e**), reproducing the well-established findings from empirical TMS studies [25]. During the cTB-TMS-40s simulation, the moving average of [*Ca*^2+^] remains predominantly in the LTD region of the Ω-function, with an overall mean [*Ca*^2+^] of 0.165*µM* (**Fig. 3e**). In the cTB-TMS-80s simulation, 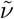 initially decreases because of LTD induced by relatively low moving average [*Ca*^2+^] during the first 40*s*, and subsequently increases monotonically during the later phase as larger [*Ca*^2+^] induces LTP (**Fig. 3e**). Notably, the change in *ν* induced by the TMS protocols is smaller than that induced by the TUS protocols, despite the higher peak [*Ca*^2+^] observed during TMS (**Fig. 3e**). This difference arises from the distinct waveform structures of TUS and TMS. TUS pulses are smoothly varying, whereas TMS pulses are discrete; consequently, TUS induces more sustained calcium responses, while TMS produces a spaced set of calcium transients that have more time to decay before the next pulse (**Fig. 3e**; see also **Fig. S2** for additional details). The simulated time series shown in **Fig. 3e** are truncated to emphasize calcium dynamics during stimulation, whereas the complete time series are provided in **Fig. S3**. These simulations demonstrate that the temporal profile of intracellular calcium, rather than its peak amplitude alone, determines the resulting plasticity outcome by governing the evolution of synaptic strength through the CaDP mechanism.

**Fig. 3d** illustrates one possible molecular interpretation of the modeled CaDP mechanism. NMDA receptors regulate intracellular calcium influx in a voltage-dependent manner through extracellular *Mg*^2+^ blockade, whereas intracellular calcium modulates AMPA receptor conductance, which is widely considered the primary mechanism underlying the expression of synaptic plasticity [54, 55]. In this interpretation, increased AMPA receptor conductance corresponds to LTP, whereas decreased AMPA receptor conductance corresponds to LTD (see **Discussion**).

### Model-based predictions for novel TUS protocols

Thus far our focus has been strictly on simulation of reported TUS and TMS experimental protocols and replication of the plasticity results in these studies. To conclude this investigation, we turned our attention to the distinctive feature of mathematical modeling in discovery science — namely as a tool for exploring novel areas of parameter space for interesting or useful properties. In doing this we sought to make strong predictions about as-yet unseen protocols that may (if confirmed) further strengthen confidence in the validity of the theory. Simulations of the CaDP-endowed circuit model under varying TUS parameters not yet observed experimentally, including pulse amplitude (*A*_*u*_), sonication duration, PRF, and duty cycle are shown in (**Fig. 4**). Unless otherwise specified, the default values of sonication duration, PRF, and duty cycle in **Fig. 4**, were set to 80*s*, 5*Hz*, and 10%, respectively [29]. The default TUS pulse amplitude (*A*_*u*_) is indicated by the red star in **Fig. 4a**, and was held fixed in **Fig. 4b-e**. This amplitude was determined using the equivalent-energy principle (**Eq. 1**) to match a standard set of TMS stimulation parameters known to induce plasticity [33, 41]. The plasticity prediction corresponding to this energy-constrained *A*_*u*_ shows strong agreement with the empirical findings reported by Zeng et al. [29] using the same default TUS configuration (80*s*, 5*Hz*, and 10%; purple star in **Fig. 4a**). **Fig. 4a** further shows that the model predicts the existence of an effective range of TUS amplitudes that enhances cortical excitability. Beyond this range, the predicted plasticity effect decreases as TUS amplitude continues to increase. In addition, **Fig. 4e** demonstrates a pronounced decreasing trend in TUS-induced excitability as the duty cycle increases from 5%. Under the condition in **Fig. 4e**, increasing the duty cycle corresponds to a more intensive stimulation protocol, because the same total stimulation energy (i.e., total TUS-ON duration) is delivered over a shorter period of time. Therefore, the model predicts that more intensive TUS protocols – corresponding to greater energy delivery per unit time – may reduce stimulation-induced plasticity effects. This prediction is consistent with empirical observations reported in TMS studies [56, 57].

**Figure 4:**
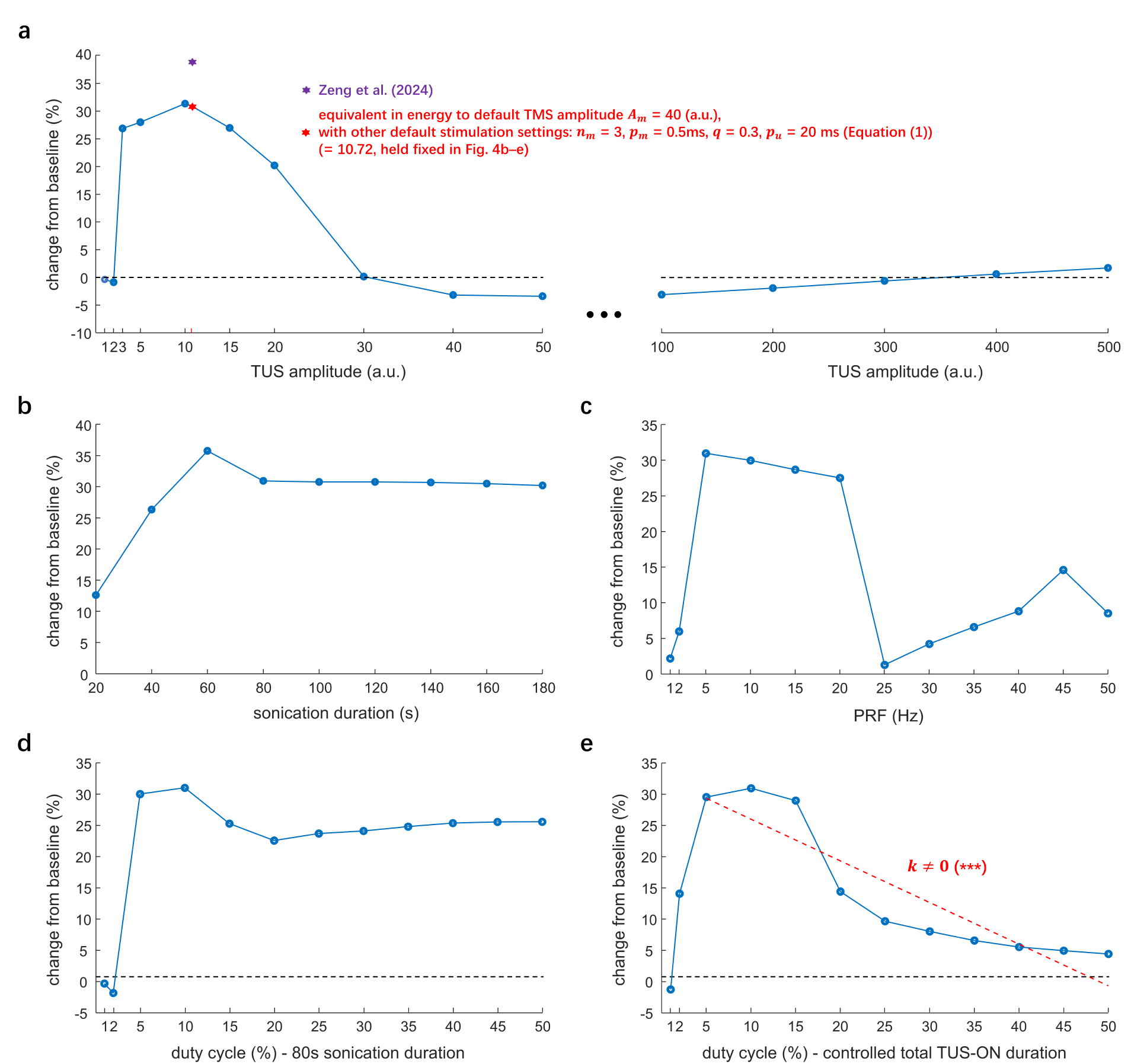
Unified TUS-TMS plasticity model predictions for unseen TUS protocols. We simulate the model in response to varying TUS parameters, including amplitude (phenomenological), sonication duration, pulse repetition frequency, and duty cycle. The TUS-induced plasticity effect is represented by the percentage change in synaptic weight (*ν*_*ee*_) relative to its pre-TUS baseline value. Unless otherwise specified, the default values of sonication duration, PRF and duty cycle are set to 80*s*, 5*Hz*, and 10%, respectively. The default value of TUS amplitude is indicated in **a)** by the red star. **a)** *TUS-induced plasticity effect in response to varying phenomenological amplitudes*. The red star indicates the plasticity effect corresponding to the TUS amplitude value (= 10.72) obtained from the equivalent-energy principle (**Eq. 1**) using the default stimulation settings. The purple star represents the plasticity effect reported by Zeng et al. (2024) under the same default TUS configuration (80*s* sonication duration, 5*Hz* PRF and 10% duty cycle). Remaining panels show TUS-induced plasticity effects in response to varying sonication durations (**b**), PRFs (**c**), and duty cycles (**d** and **e**); TUS amplitude is maintained at 10.72 (red star in **a)**) in these panels. Panel **e)** shows TUS-induced plasticity effects in response to varying duty cycles while controlling for total TUS-ON duration. For each duty cycle, the sonication duration is adjusted such that the total TUS-ON time remains constant. A regression line is fitted starting from 5% duty cycle, and the fitted slope is significantly different from 0 (two-sided t-test, *p* = 2.5 *×* 10^−4^ *<* 0.001), demonstrating that the plasticity effect decreases as duty cycle increases when the total TUS-ON duration is fixed.

In **Fig. 4b**, our model predicts that the plasticity effect generally increases and saturates as the sonication duration increases. Consistent with this prediction, the empirical results reported by Zeng et al. [29] showed increasing cortical excitability with increasing sonication duration, while the difference in plasticity between 80*s* and 120*s* was smaller than the difference between 40*s* and 80*s*. Regarding variations in PRF, the model predicts a nonlinear relationship, with the plasticity effect increasing at low PRFs, reaching a maximum at approximately 5*Hz*, and generally decreasing at higher PRFs (**Fig. 4c**). This prediction is consistent with the nonlinear trend observed experimentally by Zeng et al. [29], who similarly reported the strongest plasticity effect at 5*Hz* PRF. Duty cycle also exhibits a nonlinear relationship with the predicted plasticity response when sonication duration is held constant (**Fig. 4d**). Plasticity increases at relatively low duty cycles, reaching its largest values around 10%, whereas further increases in duty cycle do not necessarily enhance plasticity and can instead reduce the response (**Fig. 4d**). This nonlinear dependence is consistent with the findings of Zeng et al., who reported that, under the same sonication duration (80*s*) and PRF (5*Hz*), increasing duty cycle from 5% to 10% significantly enhanced plasticity, whereas a further increase from 10% to 15% produced no significant additional effect [29]. In addition, another study involving systematic TUS parameter exploration reported that varying the duty cycle did not significantly affect TUS-induced plasticity in M1 [58]. Taken together, these observations suggest that duty cycle does not have a simple monotonic relationship with plasticity.

A different pattern emerges when the total TUS-ON duration - representing total delivered energy - rather than total protocol duration, is held constant **Fig. 4e**. The plasticity response initially increases at low duty cycles and reaches a maximum around 5 − 15%; from 5% onward, however, increasing duty cycle is associated with a significant overall decrease in the predicted plasticity response (*k*≠ 0, *p <* 0.001) (**Fig. 4e**). Because total TUS-ON duration is fixed, increasing duty cycle delivers the same cumulative sonication over a progressively shorter total protocol duration. The model therefore predicts that concentrating stimulation into a shorter time does not necessarily enhance plasticity and may instead reduce it. Conversely, distributing the same cumulative sonication over a longer period may favor stronger plasticity responses. This prediction motivates future investigation of temporally distributed TUS protocols, potentially including stimulation blocks separated by intervals, which may additionally help limit cumulative tissue heating and treatment intolerance [59].

## Discussion

In this work, we developed a computational model of neuronal plasticity induced by low-intensity ultrasonic stimulation in the human brain. The model describes a corticothalamic circuit endowed with a calcium, NMDA, and AMPA-based synaptic weight update mechanism, based on existing models of the plasticity-inducing effect of repetitive TMS. Our investigations with this model led us to also develop a novel mathematical approach for defining equivalent TMS and TUS parameter configurations, in terms of the energy delivered. This allowed us to posit an equivalent-energy principle, and led to our key overall result that the same model, with exactly the same parameters, was able to capture a wide range of empirically-reported plasticity effects from both TMS and TUS studies. In the empirical data, neuronal plasticity is represented as changes in MEP amplitude, a widely used biomarker of cortical excitability. In addition, we simulated the model under varying TUS parameter configurations and obtained predictions that were broadly consistent with experimental observations. Our results thus provide a framework for mechanistic physiological interpretation of observed plasticity effects, for optimization of candidate TUS parameters to improve therapeutic efficacy, and for theory-driven discovery of novel protocol designs.

### Physical mechanisms of TUS

Plasticity-inducing neuromodulatory interventions such as TUS, TMS, and transcranial electrical stimulation (TES) [60] are, we suggest, best understood comprising two distinct (but interacting) mechanisms of action. The *primary* mechanism of action concerns the immediate physical and biophysical effects (acoustic waves, magnetic fields, electrical currents, etc), which are specific to each modality and require detailed characterization at both macro- and micro-spatial and temporal scales. These processes culminate in a near real-time modulation of neural activity - typically quantified in terms of perturbations to membrane potentials and/or firing rates. The *secondary* mechanism of action concerns how the resulting temporally-patterned stimulation-induced neural activity engages intrinsic cellular and circuit-level plasticity mechanisms to establish long-lasting effects. Unlike the primary mechanism of action, the secondary mechanism of action is much more spread-out in time, is heavily dependent on both fast and slow aspects of the stimulation waveform structure (spanning seconds to tens of minutes), is impacted by both present and past system state, and many other factors. Also unlike the primary mechanism, the secondary mechanism is common (albeit differentially engaged) across all physical stimulation modalities - offering a powerful point of convergence across the rapidly growing suite of acoustic, magnetic, electrical and optically-based noninvasive neuromodulation technologies.

In the present work our focus is on the secondary mechanism action, and its mapping between TUS and TMS. Accordingly, the neuronal effect of TUS is represented as a sub-kilohertz sinusoidal input, with oscillation frequencies within the range to which neuronal networks can respond [61, 62], without explicitly modeling the preceding physical processes through which the ultrasonic waveform alters neuronal membrane potential and firing. Some understanding of the primary mechanism is nevertheless essential for developing models and theories of the secondary mechanism.

TUS can influence neurons and neuronal networks through multiple physical mechanisms, including thermal effects [12], intramembrane cavitation caused by gas-bubble formation [11], pressure-wave propagation [10], and membrane vibration [7]. In addition, TUS waveforms propagate through the skull, where they can generate heating effects [63] and undergo sound-pressure attenuation depending on skull properties and TUS configurations [64]. For neuromodulation, a key question is how these physical processes translate into changes in neuronal membrane potential and firing activity, which consititute the neuronal-level inputs relevant to network dynamics [7, 65, 66]. TUS may influence membrane potential and the associated spiking activity through several possible mechanisms: i) TUS-induced thermal effects may facilitate membrane movements that trigger action potentials [7]; ii) TUS may induce the formation of gas bubbles whose oscillations produce sonoporation of the lipid membrane and mechanical stretch of ion channels, thereby altering the membrane permeability and affecting current flow including calcium influx [65, 67]; iii) TUS-generated sound pressure may mechanically deform the membrane through flexoelectric effects, whereby membrane deformation induces changes in membrane potential [7], possibly through modulation of mechanosensitive calcium channels [68].

Despite their different biophysical origins, these candidate primary mechanisms become relevant to the present neural-system model insofar as they alter neuronal membrane potential and consequently firing activity. At this level, TUS shares a common functional consequence with other neuromodulation techniques, such as TMS and deep brain stimulation [66], which also alter neuronal membrane potential and firing through electrically mediated mechanisms. In our framework, these modality-specific primary mechanisms converge onto neuronal dynamics, after which the common secondary CaDP mechanism links temporally patterned activity to lasting synaptic plasticity. Nevertheless, TUS transfers energy to neural tissues through fundamentally different physical processes from electromagnetic stimulation, and these modality-specific processes are not explicitly represented in the present model. Incorporating such mechanisms into future models could provide a more physiologically grounded mapping from the applied TUS waveform to membrane-potential and firing-rate dynamics, and may help account for the remaining discrepancies between simulated and empirical TUS responses (**Fig. 2a**).

### Possible interpretations and implications of the equivalent-energy principle

The intensity of energy delivered by neuromodulation devices is an important determinant of stimulation-induced plasticity [56, 29, 69]. The delivered energy must remain within an appropriate range, as insufficient stimulation may fail to produce measurable effects, whereas excessive stimulation might induce adverse side effects [70]. Moreover, increasing stimulation intensity does not necessarily enhance plasticity-related changes in cortical excitability and may instead attenuate or even reverse them [56, 57]. Stimulation intensity must therefore be considered together with parameters that determine the temporal pattern of energy delivery, including amplitude [71], frequency [69], pulse width [72], and duty cycle [29].

Our equivalent-energy principle was motivated by a parsimonious, Occam’s-razor approach to relating TUS and TMS. Rather than introducing separate plasticity mechanisms or additional modality-specific neuronal parameters, we asked whether the simplest correspondence – matching the energy associated with their stimulation waveforms – could provide a useful mapping between the two modalities while retaining the same underlying neural circuit and CaDP mechanisms. Considerable efforts have been devoted to exploring the TMS parameter spaces for improving therapeutic plasticity effects in neurological disorders [73–75], whereas systematic exploration of TUS parameter configurations, particularly for clinical applications, remains relatively limited. The equivalent-energy principle (**Eq. 1**) provides an algebraic relationship between TMS parameters, including amplitude, frequency, and pulse width, and the corresponding TUS parameters, including amplitude, oscillation, and duty cycle. It therefore provides a simple means of using established TMS protocols as reference points for exploring the less well-characterized TUS parameter space.

The results in **Fig. 2** show that, when TUS amplitude is calibrated using the equivalent-energy relationship, the same neural circuit and CaDP model can reproduce plasticity effects broadly consistent with empirical observations across both TUS and TMS protocols. Importantly, this correspondence is not strongly dependent on the precise oscillatory form assumed for the effective TUS input. The equivalent-energy calculation depends on the ratio *q* between the oscillatory and mean amplitudes of the sinusoidal TUS waveform (**Eq. 1**); however, numerical tests using a non-oscillatory box-shaped TUS input produced plasticity outcomes similar to those reported here, with *q* = 0.3 providing only a modest improvement in agreement with the empirical data. Furthermore, under the assumptions of the equivalent-energy derivation, the TUS oscillation frequency does not enter the resulting energy relationship. Together, these observations suggest that the model predictions depend primarily on the effective stimulation magnitude and broader temporal pattern of energy delivery, rather than on the precise shape or frequency of the fast oscillatory component.

This robustness provides one interpretation of why such a simple equivalent-energy mapping can be informative despite the substantial physical differences between TUS and TMS. At the level of neuronal plasticity represented in the present model, detailed features of the high-frequency TUS waveform may be less important than the effective neuronal drive and its temporal organization. This does not imply that the underlying ultrasonic carrier frequency or waveform is physiologically unimportant; these features can strongly influence ultrasound propagation and interaction with neural tissue through physical mechanisms that are outside the scope of the present model. Rather, our results indicate that once the physical effects of TUS are represented as an effective neuronal input, the predicted CaDP response is relatively robust to the precise oscillatory representation of that input.

Nevertheless, broader differences in temporal waveform structure between TUS and TMS remain important in the model. Specifically, cTB-TMS produced different plasticity directions (LTP or LTD) depending on stimulation duration, whereas cTB-TUS consistently induced LTP, with stimulation duration primarily affecting the magnitude of potentiation (**Fig. 2a**). One possible explanation suggested by the model is that the discrete, high-amplitude pulses of TMS generate more abrupt intracellular calcium transients that cross different CaDP regimes as stimulation progresses, whereas the more continuous temporal profile of TUS produces smoother calcium dynamics and consequently more gradual modulation of plasticity. Beyond this model-based explanation, the observed differences may also reflect the fundamentally different physical mechanisms through which electromagnetic and acoustic stimulation interact with neural tissue, which are not explicitly represented in the present model. Future modeling incorporating ultrasound propagation, tissue mechanics and their effects on neuronal membrane dynamics could provide a more complete account of how these modality-specific physical mechanisms shape neuronal activity and plasticity.

### Cellular substrates of synaptic plasticity

At the molecular level, the CaDP mechanism is reflected in the dynamics of membrane conductance mediated by ion-channel gating [76, 77]. In particular, the NMDA receptor is the primary mediator of synaptic plasticity induction, including LTD or LTP through its regulation of intracellular calcium concentration [*Ca*^2+^] [55, 54]. NMDA receptor conductance, which mediates calcium influx, is voltage dependent [55] (**Fig. 3b, Eq. 7**), exhibits relatively slow kinetics [54] (represented by the large time constant in **Eq. 13**), and is modulated by extracellular magnesium (*Mg*^2+^) blockade [78] (**Fig. 3d, Eq. 10**). Intracellular [*Ca*^2+^] can regulate synaptic plasticity by modulating the dynamics of the AMPA receptors, which primarily mediate sodium (*Na*^+^) conductance [76, 55]. AMPA receptor conductance is considered a principal mechanism underlying the expression of synaptic plasticity [54, 55]. In contrast to NMDA receptors, AMPA receptors exhibit relatively fast [54] dynamics, and their contribution to synaptic transmission can be modified through changes in receptor number/density on the postsynaptic membrane [76] and the single-channel conductance [54]. Both processes can be regulated by intracellular [*Ca*^2+^] through NMDA receptor-mediated signaling pathways [76] (**Fig. 3d**). A pharmacological study showed that the modulatory effect of cTB-TUS is significantly attenuated by the administration of a *Na*^+^ channel blocker, *Ca*^2+^ channel blocker, or NMDA receptor antagonist [79], indicating that sodium and calcium signaling, together with NMDA receptor activity, play critical roles in TUS-induced plasticity.

The molecular basis underlying the calcium-dependent synaptic weight regulation, represented in our model principally by the Ω-function abstraction and the calcium dynamics of **Eqs. 7-17**, is believed to capture a differential activation of downstream calcium-sensitive signaling pathways. Specifically, the inverted-Ω-shape dependence of the calcium-control function on intracellular [*Ca*^2+^] reflects differences in how two opposing *Ca*^2+^-binding effectors, calcium/calmodulin-dependent protein kinase II (CaMKII) and calcineurin (CaN), respond to intracellular calcium. CaMKII activation requires *Ca*^2+^ levels high enough to drive cooperative T286 autophosphorylation – a threshold preferentially crossed during the brief, high-amplitude *Ca*^2+^ transients characteristic of LTP-inducing stimulation [80, 81]; CaN activates more readily at the lower, sustained *Ca*^2+^ elevations that drive LTD [82]. Both pathways read out at GluA1 phosphorylation, but in opposite directions: CaMKII phosphorylation at S831 increases AMPA recptor single-channel conductance and supports synaptic potentiation, while CaMKII phosphorylation at S567 and calcineurin-driven dephosphorylation reduce synaptic AMPA receptor content [83, 82]. Accordingly, the calcium-control-Ω-function (**Eq. 17**) can be viewed as a phenomenological approximation of this kinase-phosphatase balance [40]. Together, these cellular mechanisms provide a biological interpretation of the mathematical abstractions underlying CaDP in our model. Although the model does not explicitly simulate individual receptor, kinase, or phosphatase dynamics, NMDA receptor-mediated calcium dynamics and the Ω-function capture their aggregate effects on the induction and direction of synaptic plasticity. This provides a mechanistic basis for linking stimulation-induced neuronal activity and calcium dynamics to the LTP- and LTD-like outcomes predicted by the model.

### Future model extensions

The corticothalamic circuit and CaDP models used in this work were adapted from previous models investigating TMS-induced plasticity [41, 33]. These models can be further adjusted, refined, or extended depending on the specific research questions being addressed, including the spatial or temporal scales of interest and the particular plasticity mechanisms under investigation. For example, short-term synaptic plasticity (STP) – which refers to transient synaptic changes lasting from milliseconds to minutes following external stimulation [84] – may also play an important role in both TMS and TUS mechanisms. However, STP was not incorporated into the present model because accurate STP modeling requires more detailed representations of synaptic physiology, including neurotransmitter release and recovery dynamics, and receptor-specific mechanisms [85]. Beyond the structural refinements of the model, the model parameters themselves may also be sharpened based on additional physiological evidence. For instance, the time constants governing the CaDP dynamics – including calcium dynamics (**Eq. 7**) and NMDA receptor-conductance-mediated metaplasticity dynamics (**Eq. 13**) – could be further validated and refined. Adjustments to these parameters may help account for quantitative differences in the PRF-dependent plasticity response between the model predictions and empirical observations (**Fig. 2c**). In addition, the model could be extended by incorporating more detailed molecular mechanisms underlying the expression of plasticity, particularly the process through which intracellular [*Ca*^2+^] regulates plasticity via modulation of AMPA receptor conductance [76, 40]. Although the plasticity-induction-Ω-function (**Eq. 17**) has been used as a phenomenological representation of the AMPA receptor conductance change in response to intracellular [*Ca*^2+^] [40], a more explicit characterization of AMPA receptor dynamics, together with NMDA receptor dynamics [86], could provide deeper physiological insight into the mechanisms represented by our model.

### More empirical datasets for future model validations and developments

TUS neuromodulation remains an emerging technology, and there is currently a limited number of empirical datasets characterizing neural responses to different TUS configurations and stimulation patterns. In **Fig. 4**, we simulated the model under varying TUS parameters, and these predictions could be further validated or refined using future experimental datasets, which will be critical for continued model development. For example, in **Fig. 4e**, when the total TUS-ON duration - the delivered energy - is held constant, the model predicts an overall decrease in the magnitude of induced plasticity as duty cycle increases from 5% onward. One possible explanation of this attenuation is short-term synaptic depression induced by intensive external stimulation, similar to effects observed during high-frequency DBS, where persistent simulation can lead to synaptic depletion or reduction in intracellular calcium availability [87]. The prediction shown in **Fig. 4e** is partially consistent with existing experimental findings. Chen et al. (2024) [88] reported inhibitory effects induced by TUS with a 30% duty cycle (pulse width = 0.3*ms*, PRF = 1*kHz*, sonication duration = 30*s*, corresponding to a total TUS-ON duration of 9*s*). In contrast, Zeng et al. (2022) [89] reported excitatory effects using cTB-TUS with a 10% duty cycle (pulse width = 20*ms*, PRF = 5*Hz*, sonication duration = 80*s*, corresponding to a total TUS-ON time of 8*s*). Although the total TUS-ON durations in the two protocols were similar (9*s* versus 8*s*), the protocol used by Zeng et al. (2022) produced stronger excitatory effects, qualitatively compatible with our prediction that, beyond the low-duty-cycle range, increasing duty cycle can attenuate TUS-induced plasticity (**Fig. 4e**). However, an important difference between these two studies is that Chen et al. (2024) used a high PRF (1*kHz*), whereas Zeng et al. (2022) applied theta-frequency (5*Hz*) TUS pulses. High-PRF TUS has previously been associated with suppressive effects in M1 [58]. Although the present model does not reproduce a transition to inhibitory plasticity over the PRF range examined, it predicts a nonlinear PRF dependence, with maximal potentiation at approximately 5*Hz* and generally weaker plasticity at higher PRFs (**Fig. 4c**). The mechanisms underlying experimentally observed suppression at substantially higher PRFs therefore remain to be investigated in future extensions of the model. In contrast to Zeng et al. (2022), Chen et al. (2026) reported that cTB-TUS with a doubled duty cycle (20%) and a halved sonication duration (40*s*) produced inhibitory, rather than facilitatory plasticity effects [90]. Although our model does not predict a complete reversal from LTP to LTD under these conditions, it similarly predicts that, for the same delivered energy, increasing the duty cycle from 10% to 20% substantially attenuates LTP (**Fig. 4e**). New experimental datasets specifically examining plasticity responses across varying duty cycles while controlling for total TUS-ON duration (as in **Fig. 4e**) would be particularly valuable for elucidating the mechanisms through which duty cycle influences TUS-induced plasticity.

TUS data collected using intermittent theta-burst paradigms – allowing several seconds of TUS-OFF intervals between stimulation periods – may also provide an important direction for future exploration. This possibility is motivated by the strong clinical efficacy of the intermittent TMS protocol iTB-TMS-200s (also referred to as iTBS600; **Fig. 2**) in disorders such as depression [23], stroke [17], and Parkinson’s disease [24]. Beyond facilitating model validation and refinement using new datasets, existing TUS datasets obtained using established stimulation protocols also raise new mechanistic questions [28, 88]. For example, Xia et al. (2024) investigated the remote contralateral effects of cTB-TUS and found that left-M1 cTB-TUS reduced excitability in right M1 [28]. In addition, Chen et al. (2024) performed concurrent and sequential stimulation experiments of TUS and TMS, and demonstrated that TUS and TMS may interact in modulating cortical excitability [88]. Our model implements a BCM-like metaplasticity mechanism in which previous stimulation modifies NMDA receptor efficacy, thereby altering subsequent calcium dynamics and shifting the conditions under which LTP or LTD is induced. This provides a mechanistic framework potentially capable of explaining experimental priming effects, such as the prolonged plasticity effects observed when cTB-TMS precedes cTB-TUS [91].

## Methods

### An equivalent-energy principle for determining TUS vs. TMS input scaling

The two noninvasive brain stimulation techniques considered in this study (i.e., TUS and TMS) were represented as temporally structured inputs to the neural populations. For TUS, the effect of the megahertz sonication waveforms generated by the ultrasound transducers [29, 92] is represented in the model as a sub-kilohertz sinusoidal stimulating input, within the frequency range to which neuronal systems can respond [61, 62]. The resulting TUS input is represented as sinusoidal bursts delivered at a pulse repetition frequency (PRF), defined as the reciprocal of the inter-burst interval, with each burst oscillating at 500 Hz [93] as shown in **Fig. 1**. Moreover, the TMS input is represented as rectangular pulses delivered in bursts, with burst repetition frequency likewise defined as the reciprocal of the inter-burst interval [33]. The TUS model parameters are then constrained using previously established TMS model parameters that have successfully reproduced plasticity effects consistent with empirical TMS data [41, 33].

For the purpose of determining TUS model parameters from established TMS configurations, an equivalent-energy principle was developed to link the TUS and TMS waveforms such that the same amount of energy is delivered to the neural populations by both noninvasive neuromodulation techniques. This principle as portrayed in **Fig. 1** is defined as

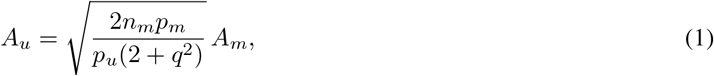

where *A*_*u*_ is the mean amplitude of TUS sinusoidal waveform, *A*_*m*_ is the amplitude of a TMS rectangular pulse, *n*_*m*_ is the number of pulses within each TMS burst, *p*_*m*_ is the width of a TMS pulse, *p*_*u*_ is the width of a TUS burst, and *q* is the ratio of the TUS oscillation amplitude to the mean amplitude *A*_*u*_. Furthermore, the mathematical derivations of this principle are provided in the **Supplementary Methods** section. Therefore, this principle provides the basis to derive TUS parameters by leveraging the extensive literature on TMS configurations [25, 43].

In this study, the default TMS configurations were set as *A*_*m*_ = 40 (a.u.), *n*_*m*_ = 3 ([25]), and *p*_*m*_ = 0.5 ms ([33]). These values remained unchanged throughout the study and produced plasticity effects consistent with empirical TMS observations. For the default TUS configuration, *q* was fixed at 0.3, and the remaining TUS parameters were determined to satisfy the equivalent-energy principle defined in **Eq. 1**. The resulting correspondence between TUS and TMS parameters was maintained across all simulations and used to compare the model-predicted plasticity effects with empirical MEP data obtained from both TUS and TMS experiments.

### Neurophysiological model of corticothalamic circuits

To model calcium-dependent synaptic plasticity effects induced by TUS and TMS, the corticothalamic circuit used here is represented by a mesoscale neural population model [41, 33]. The formalisms in this model describe states and parameters relevant to corticothalamic neural populations, denoted by *a, b* ∈ {*e, i, r, s*} (i.e., excitatory, inhibitory, relay, and reticular) and linked through distinct combinations of synaptic connections. Within this circuit, external stimulation and synaptic inputs contribute to the mean soma potential of each population.

The mean soma potential of each neural population at time *t* is defined as

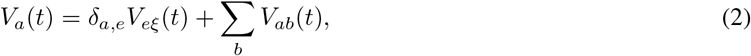

where *V*_*a*_(*t*) denotes the mean soma potential of population *a, V*_*eξ*_(*t*) represents the external stimulus produced by the TUS or TMS protocols, and *V*_*ab*_(*t*) represents the contribution arising from synaptic activity transmitted by population *b*. The Kronecker delta function *δ*_*a,e*_ equals to unity when *a* = *e* and zero otherwise, thereby restricting the external stimulation term to the cortical excitatory population. Thus, the mean soma potential combines the direct effect of stimulation with the cumulative synaptic contributions from the connected corticothalamic populations.

For both synaptic activity and external stimulation at time *t*, the corresponding contributions to the mean soma potential are generated through second-order synaptodendritic low-pass filtering, defined as

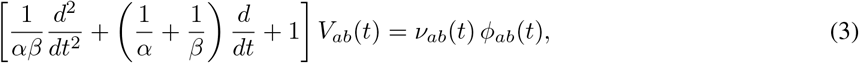

and

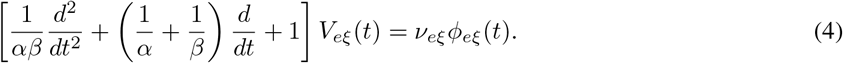

In **Eq. 3**, *ν*_*ab*_(*t*) denotes the mean synaptic coupling strength from population *b* to *a*, and *ϕ*_*ab*_(*t*) represents the propagated firing-rate activity received by population *a* from population *b*. Moreover, in **Eq. 4**, *ν*_*eξ*_ scales the strength of the external stimulus, and *ϕ*_*eξ*_(*t*) denotes the TUS and TMS waveform activity. The parameters *α* and *β* in both **Eqs. 3 and 4** represent the mean decay and rise rate of the synaptodendritic response [34]. Therefore, **Eqs. 3 and 4** transform the synaptic and stimulation inputs into their respective contributions, *V*_*ab*_(*t*) and *V*_*eξ*_(*t*), to the mean soma potential.

Following the integration of the synaptic and stimulation-induced contributions, the mean soma potential of population *a* is converted into the mean firing rate of each neural population through a sigmoidal response function which is expressed as

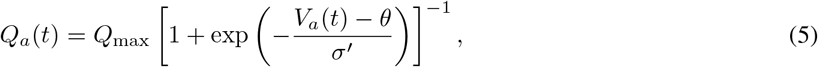

where *Q*_*a*_(*t*) is the mean firing rate of the population *a, Q*_max_ is the maximal firing rate, *θ* is the mean firing threshold, and *σ*^*′*^ characterizes the scaling factor associated with the voltage standard deviation [94]. The resulting firing activity from population *b* is subsequently transmitted along the corresponding axonal pathway according to

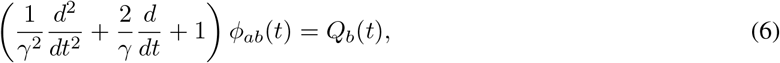

such that

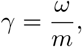

where *m* denotes the spatial range of axonal propagation, *ω* is the axonal propagation velocity, and *γ* characterizes the temporal damping of the propagated activity [95]. Together, these dynamics complete the recurrent flow of activity through the corticothalamic circuit.

### Calcium-dependent plasticity (CaDP) model

Postsynaptic intracellular calcium concentration is the key factor governing plasticity effects, including long-term potentiation (LTP) and long-term depression (LTD) [32, 38]. Moderate postsynaptic calcium levels induce LTD, whereas high calcium concentrations trigger LTP. This calcium-control hypothesis for bidirectional plasticity effects is well established and supported by empirical evidence [38, 34]. Specifically, synaptic connectivity strength is modulated by postsynaptic calcium concentration through a nonlinear, inverted-Ω-shaped relationship (**Fig. 1**) [41, 40]. Postsynaptic calcium concentration is regulated by the dynamics of the N-methyl-D-aspartate (NMDA) receptor, which receives glutamate neurotransmitters and serves as the primary pathway for intracellular calcium influx [41, 32]. Variations in NMDA conductance that influence calcium dynamics are further characterized as a metaplasticity mechanism [41, 33], originating from the Bienenstock-Cooper-Munro (BCM) theory of plasticity [96]. This theory suggests that variability in stimulation can induce cascading physiological changes via a lower-order mechanism of LTD and LTP to mediate their subsequent effects and functional outcome.

In this work, calcium-dependent plasticity (CaDP) was modeled for each synaptic connection, with intracellular calcium concentration [*Ca*^2+^] abbreviated as *Ca* in the following equations:

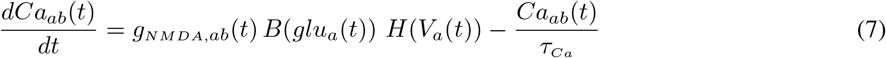

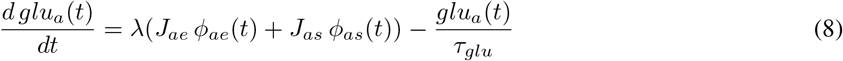

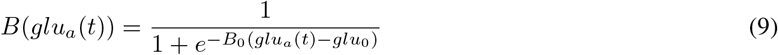

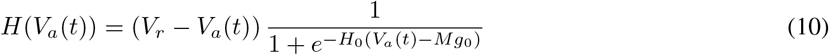

In **Eq. 7**, *Ca*_*ab*_ denotes the calcium dynamics induced by the synaptic connection *b* → *a* (*a, b* ∈ {*e, i, r, s*}). The parameter *τ*_*Ca*_ is the calcium decay time constant, 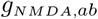 represents the time-varying and history-dependent NMDA conductance (i.e., metaplasticity, specified later), *glu*_*a*_ denotes the glutamate concentration dynamics in neuronal population *a* (**Eq. 8**), *B*(*glu*_*a*_) characterizes glutamate-binding dynamics (**Eq. 9**), and *H*(*V*_*a*_) represents the voltage dependence of NMDA conductance (**Eq. 10**). In **Eq. 8**, *λ* is a scaling constant representing the average amount of glutamate released per-spike. The variables *J*_*ae*_ and *J*_*as*_ equal 1 if the *e* → *a* and *s* → *a* connections exist, respectively (and 0 otherwise). The terms *ϕ*_*ae*_ and *ϕ*_*as*_ denote the axonally propagated and damped firing-rate activity (**Eq. 6**), while *τ*_*glu*_ is the decay time constant. Glutamate release is assumed to originate from excitatory projections arising from neuronal populations *e* or *s* [34]. In **Eq. 9**, *B*_0_ is a scaling constant for the glutamate-binding strength, and *glu*_0_ is the threshold for glutamate binding. In **Eq. 10**, *V*_*r*_ is a calibrated resting soma potential for the neural model [34], *H*_0_ is a scaling constant, and *Mg*_0_ is a constant that determines the strength of the extracellular *Mg*^2+^ blockade of NMDA-mediated calcium conductance.

The plasticity effects induced by calcium dynamics are represented by changes in synaptic connectivity strength *ν*_*ab*_:

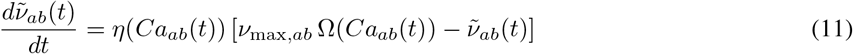

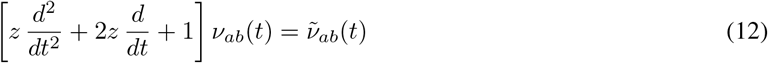

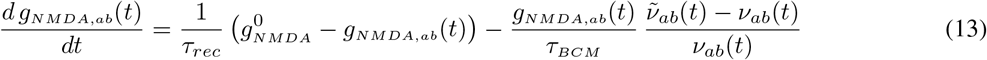

In **Eq. 11**, 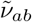 denotes the instantaneous synaptic strength induced by stimulation [41], and *ν*_max,*ab*_ is the maximal strength of the *b* → *a* connection. The functions *η*(*Ca*_*ab*_) and Ω(*Ca*_*ab*_) are key calcium-control functions governing CaDP [34], and are specified later. In **Eq. 12**, the instantaneous synaptic strength 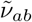 passes through a low-pass filter to generate the final stabilized synaptic strength *ν*_*ab*_, where *z* is a timescale over which the underlying plasticity processes are expressed as stabilized changes in synaptic strength. The difference between the stabilized and instantaneous strengths, 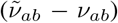, drives the metaplasticity mechanism, which is characterized by the dynamics of NMDA conductance *g*_*NMDA,ab*_ in **Eq. 13**. Here, *τ*_*rec*_ is the recovery time constant for *g*_*NMDA,ab*_ to return to its baseline value 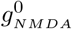, and *τ*_*BCM*_ is the characteristic timescale of BCM metaplasticity [96].

The calcium-control functions governing CaDP-induced LTP and LTD are specified as follows [34, 33, 40]:

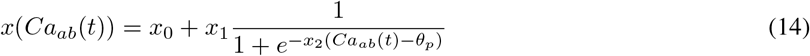

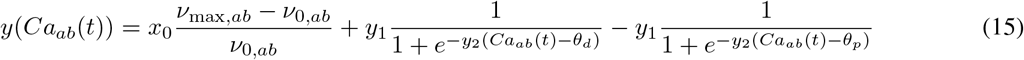

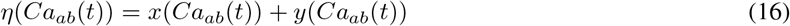

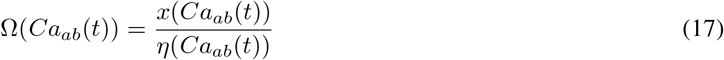

where *x* is the synaptic potentiation rate (**Eq. 14**), *y* is the synaptic depression rate (**Eq. 15**), *η* is the overall synaptic plasticity rate (**Eq. 16**), and Ω represents plasticity induction (**Eq. 17**). In **Eqs. 14**-**15**, *x*_0_ is a threshold parameter, *x*_1_ (resp., *y*_1_) determines peak potentiation (resp., depression) rate, *x*_2_ (resp., *y*_2_) controls the transition slope, and *θ*_*p*_ and *θ*_*d*_ are the potentiation and depression thresholds, respectively. In **Eq. 15**, *ν*_max,*ab*_ and *ν*_0,*ab*_ denote the maximal and initial strength of the *b* → *a* synaptic connection, respectively. **Eq. 17** defines the Ω-function governing plasticity induction - LTP or LTD - as a function of intracellular calcium concentration (illustrated in **Fig. 1**).

### Model-predicted plasticity and model parameterization

The TUS- (or TMS-) induced plasticity predicted by our model is represented as a change in cortical excitability, quantified as the relative change in *ν*_*ee*_ (the mean synaptic connection strength within the cortical excitatory population) with respect to its baseline value:

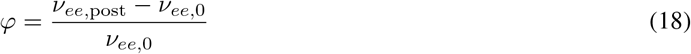

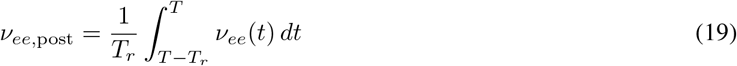

where *φ* denotes the normalized change in *ν*_*ee*_ induced by stimulation, *ν*_*ee*,0_ is the pre-stimulation strength of the *e* → *e* synaptic connection, and *ν*_*ee*,post_ is the post-stimulation synaptic strength. The quantity *ν*_*ee*,post_ is computed as the mean value of *ν*_*ee*_(*t*) over the time interval [*T* − *T*_*r*_, *T* ], where *T* is the post-stimulation time and *T*_*r*_ is the duration of the averaging interval. The model-predicted plasticity was then compared with empirical motor-evoked potential (MEP) recordings from the first dorsal interosseous (FDI) muscle on the wrist [29, 25], a widely used measure of cortical excitability in both TUS [29] and TMS [25] studies.

The parameters of the neural circuit model with CaDP are summarized in **Table S1**. All model parameters were adopted from previous studies [33, 34, 41], except for those directly associated with the Ω-function (**Eq. 17**). The Ω-function used in this work has the same functional form as that proposed by Fung and Robinson [41] (**Fig. S1**), but its parameters were re-tuned for the corticothalamic neural circuit model used here. Importantly, all parameters of the model (**Table S1**) were identical for both TUS and TMS simulations, implying that TUS and TMS are applied to the same underlying neural system. This consistency implies that the underlying neural network architecture and plasticity mechanisms in the model remain identical across different forms of external stimulation. In addition, TUS and TMS configurations are linked through the equivalent-energy principle (**Fig. 1, Eq. 1**), which was applied throughout all simulations. Our results suggest that, within computational modeling frameworks, TUS parameters may potentially be guided and constrained using existing knowledge of TMS-induced plasticity effects (see **Discussion**).

## Acknowledgments

This work was supported by grants from the Krembil Foundation, the Labbatt Foundation, the CAMH Discovery Fund, and the Fields Institute for Mathematical Sciences. Yupeng Tian was supported by the Fields Institute for Research in Mathematical Sciences through the thematic program on the Mathematics of Neuroscience during July-December 2025 (PI: John Griffiths), and by NSERC Discovery Grant RGPIN-2024-06003 (Kumar Murty), the Department of Mathematics, and the Faculty of Arts and Sciences (both of the University of Toronto).

## Data and Resource Availability

All simulations, analyses, and results described in this paper are fully reproducible with Matlab and Python code provided in the accompanying github repository at github.com/GriffithsLab/TianEtAl2026_ UnifyingTFUS-TMS-Plasticity.

## Supplementary Information

### Supplementary Table

**Table S1:** Parameters in the corticothalamic circuit model with calcium-dependent plasticity induced by TUS or TMS.

| Parameter | Description | Value | Unit | Source |
| --- | --- | --- | --- | --- |
| $\alpha$ | mean decay rate of dendrite propagation | 83.3 | $s^{-1}$ | Kadak et al. (2025) |
| $\beta$ | mean rise rate of dendrite propagation | 769.2 | $s^{-1}$ | Kadak et al. (2025) |
| $\nu_{e\xi}$ | scaling factor of the strength of external stimuli | $8.2 \times 10^{-3}$ | $mM$ | Kadak et al. (2025) |
| $Q_{max}$ | maximal firing rate | 340 | $Hz$ | Fung and Robinson (2013) |
| $\theta$ | mean firing threshold | 13 | $mV$ | Fung and Robinson (2013) |
| $\sigma'$ | scaling related to standard deviation of soma voltage | 3.8 | $mV$ | Fung and Robinson (2013) |
| $m$ | the spatial range limit of axonal propagation | 0.086 | $m$ | Kadak et al. (2025) |
| $\omega$ | axonal transmission speed | 9.976 | $ms^{-1}$ | Kadak et al. (2025) |
| $\gamma$ | axonal damping coefficient | 116 | $s^{-1}$ | Fung and Robinson (2013) |
| $\tau_{Ca}$ | calcium decay time constant | 50 | $ms$ | Fung and Robinson (2013) |
| $\lambda$ | average amount of glutamate release per-spike | 150 | $\mu Ms$ | Fung and Robinson (2013) |
| $\tau_{glu}$ | glutamate decay time constant | 30 | $ms$ | Fung and Robinson (2013) |
| $B_0$ | scaling of glutamate binding strength | $3 \times 10^4$ | $\mu M^{-1}$ | Kadak et al. (2025) |
| $glu_0$ | threshold for glutamate binding | 200 | $\mu M$ | Kadak et al. (2025) |
| $V_r$ | calibrated resting soma potential for the neural model | 195 | $mV$ | Fung and Robinson (2013) |
| $H_0$ | scaling of voltage-dependence NMDA conductance | 62 | $V^{-1}$ | Fung and Robinson (2013) |
| $Mg_0$ | impact of calcium conductance from extracellular magnesium | 45.5 | $mV$ | Fung and Robinson (2013) |
| $z$ | timescale of synaptic strength stabilization | 0.01 | $s$ | Kadak et al. (2025) |
| $\tau_{rec}$ | time constant of NMDA conductance | 1000 | $s$ | Fung and Robinson (2014) |
| $\tau_{BCM}$ | time constant of BCM metaplasticity | 7 | $s$ | Fung and Robinson (2014) |
| $x_0$ | threshold value in synaptic rates | $10^{-4}$ | $s^{-1}$ | Fung and Robinson (2013) |
| $x_1$ | related to the peak synaptic potentiation rate | 0.01 | $s^{-1}$ | this work |
| $x_2$ | scaling of change of synaptic potentiation rate | $4 \times 10^7$ | $\mu M^{-1}$ | Kadak et al. (2025) |
| $y_1$ | related to the peak synaptic depression rate | 0.05 | $s^{-1}$ | this work |
| $y_2$ | scaling of change of synaptic depression rate | $4 \times 10^7$ | $\mu M^{-1}$ | Kadak et al. (2025) |
| $\theta_d$ | depression threshold in calcium-control functions | 0.2 | $\mu M$ | this work |
| $\theta_p$ | potentiation threshold in calcium-control functions | 0.336 | $\mu M$ | this work |
| $\nu_{0,ee}$ | initial strength of $e \rightarrow e$ connection | 1.2 | $mM$ | Kadak et al. (2025) |
| $\nu_{0,ei}$ | initial strength of $i \rightarrow e$ connection | -9.8 | $mM$ | Kadak et al. (2025) |
| $\nu_{0,es}$ | initial strength of $s \rightarrow e$ connection | 9.7 | $mM$ | Kadak et al. (2025) |
| $\nu_{0,ie}$ | initial strength of $e \rightarrow i$ connection | 1.5 | $mM$ | Kadak et al. (2025) |
| $\nu_{0,ii}$ | initial strength of $i \rightarrow i$ connection | -9.6 | $mM$ | Kadak et al. (2025) |
| $\nu_{0,is}$ | initial strength of $s \rightarrow i$ connection | 9.7 | $mM$ | Kadak et al. (2025) |
| $\nu_{0,se}$ | initial strength of $e \rightarrow s$ connection | 1.1 | $mM$ | Kadak et al. (2025) |
| $\nu_{0,sr}$ | initial strength of $r \rightarrow s$ connection | 1.2 | $mM$ | Kadak et al. (2025) |
| $\nu_{0,re}$ | initial strength of $e \rightarrow r$ connection | 0.14 | $mM$ | Kadak et al. (2025) |
| $\nu_{0,rs}$ | initial strength of $s \rightarrow r$ connection | 0.058 | $mM$ | Kadak et al. (2025) |
| $\nu_{max,ee}$ | maximal strength of $e \rightarrow e$ connection | 10 | $mM$ | Kadak et al. (2025) |
| $\nu_{max,ei}$ | maximal strength of $i \rightarrow e$ connection | -100 | $mM$ | Kadak et al. (2025) |
| $\nu_{max,es}$ | maximal strength of $s \rightarrow e$ connection | 100 | $mM$ | Kadak et al. (2025) |
| $\nu_{max,ie}$ | maximal strength of $e \rightarrow i$ connection | 10 | $mM$ | Kadak et al. (2025) |
| $\nu_{max,ii}$ | maximal strength of $i \rightarrow i$ connection | -100 | $mM$ | Kadak et al. (2025) |
| $\nu_{max,is}$ | maximal strength of $s \rightarrow i$ connection | 100 | $mM$ | Kadak et al. (2025) |
| $\nu_{max,se}$ | maximal strength of $e \rightarrow s$ connection | 10 | $mM$ | Kadak et al. (2025) |
| $\nu_{max,sr}$ | maximal strength of $r \rightarrow s$ connection | -10 | $mM$ | Kadak et al. (2025) |
| $\nu_{max,re}$ | maximal strength of $e \rightarrow r$ connection | 10 | $mM$ | Kadak et al. (2025) |
| $\nu_{max,rs}$ | maximal strength of $s \rightarrow r$ connection | 10 | $mM$ | Kadak et al. (2025) |

### Supplementary Methods - derivation of Eq. 1

The equivalent-energy principle 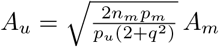 (**Eq. 1**) computes the mean amplitude *A*_*u*_ of a sinusoidal TUS burst based on the remaining TUS and TMS parameters, such that the delivered energy by the TMS and TUS waveforms is equivalent, assuming the same inter-burst-interval (IBI), whose reciprocal corresponds to the carrier frequency of the bursts in the stimulation waveforms (**Fig. 1**). The energy of a signal is defined as the integral of the squared signal amplitude over time [97]. Accordingly, the energy of a TMS burst is given by:

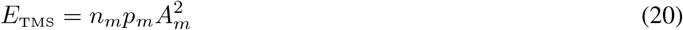

where *n*_*m*_ is the number of pulses within each TMS burst, *p*_*m*_ is the width of a TMS pulse, *A*_*m*_ is the amplitude of a rectangular TMS pulse, and *E*_TMS_ denotes the energy of one burst of TMS pulses. The energy of a TUS burst is given by:

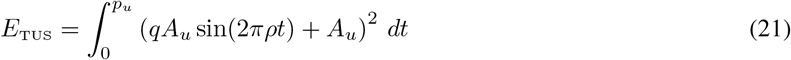

where *A*_*u*_ is the mean burst amplitude, *qA*_*u*_ is the oscillation amplitude of the sinusoidal waveform, *q* is the ratio of sinusoidal oscillation amplitude to the mean burst amplitude, *ρ* is the oscillation frequency, *p*_*u*_ is the width of a TUS burst, and *E*_TUS_ denotes the energy of a TUS burst. After algebraic simplification, **Eq. 21** can be written as:

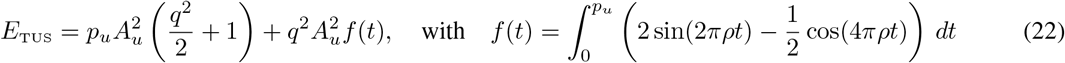

In **Eq. 22**, the sinusoidal term sin(2*πρt*) has period 1*/ρ*, while cos(4*πρt*) has period 1*/*(2*ρ*). Because *ρ* represents the oscillation frequency of the TUS burst, the burst width *p*_*u*_ can be expressed as an integer number (*N* ) of oscillation periods (1*/ρ*), together with a small residue term *ε* that is smaller than one oscillation period. The residue becomes zero when *p*_*u*_ is an exact integer multiple of 1*/ρ*. Since a TUS burst (*p*_*u*_) is expected to contain many oscillation cycles (*N* ≫ 1), it is reasonable to assume that:

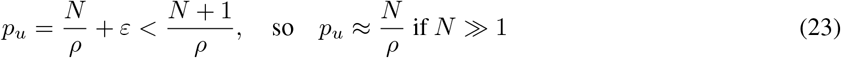

Therefore, continuing from **Eq. 22**,

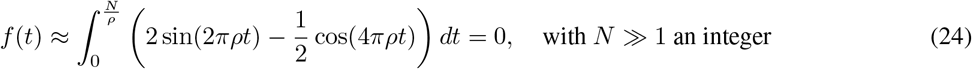

because the integrals of sin(2*πρt*) and cos(4*πρt*) over integer multiples of their respective periods are equal to zero. Combining **Eqs. 20**,**22**,**24**, the equivalent-energy relationship between TMS and TUS parameters becomes:

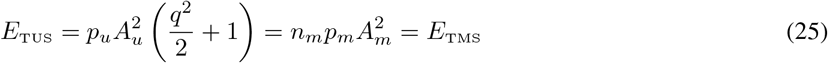

The equivalent-energy principle (**Eq. 1**) is then obtained directly from **Eq. 25**.

### Supplementary Figures

**Figure S1:**
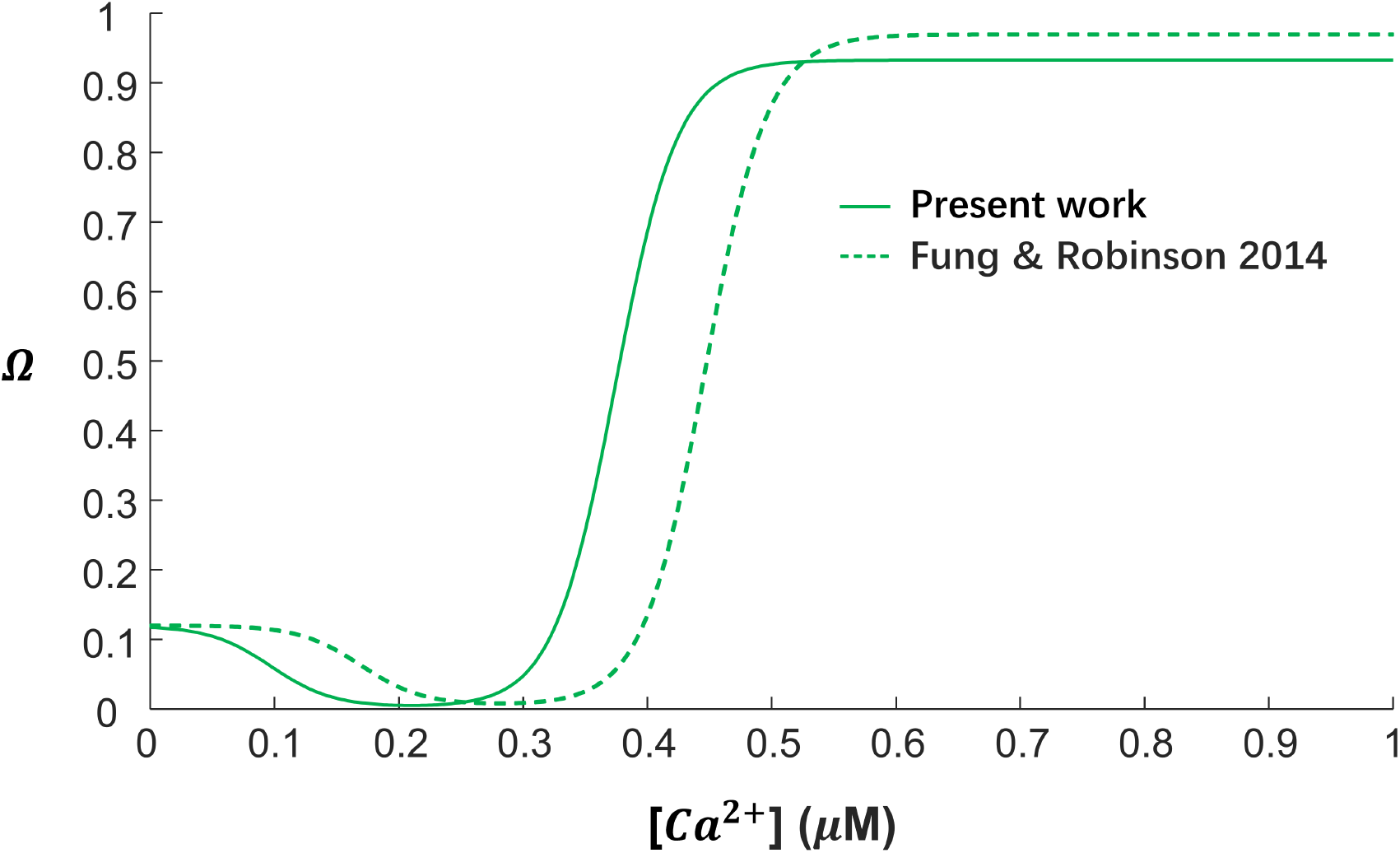
Ω-functions used in this work (Eq. 17) and Fung & Robinson (2014)

**Figure S2:**
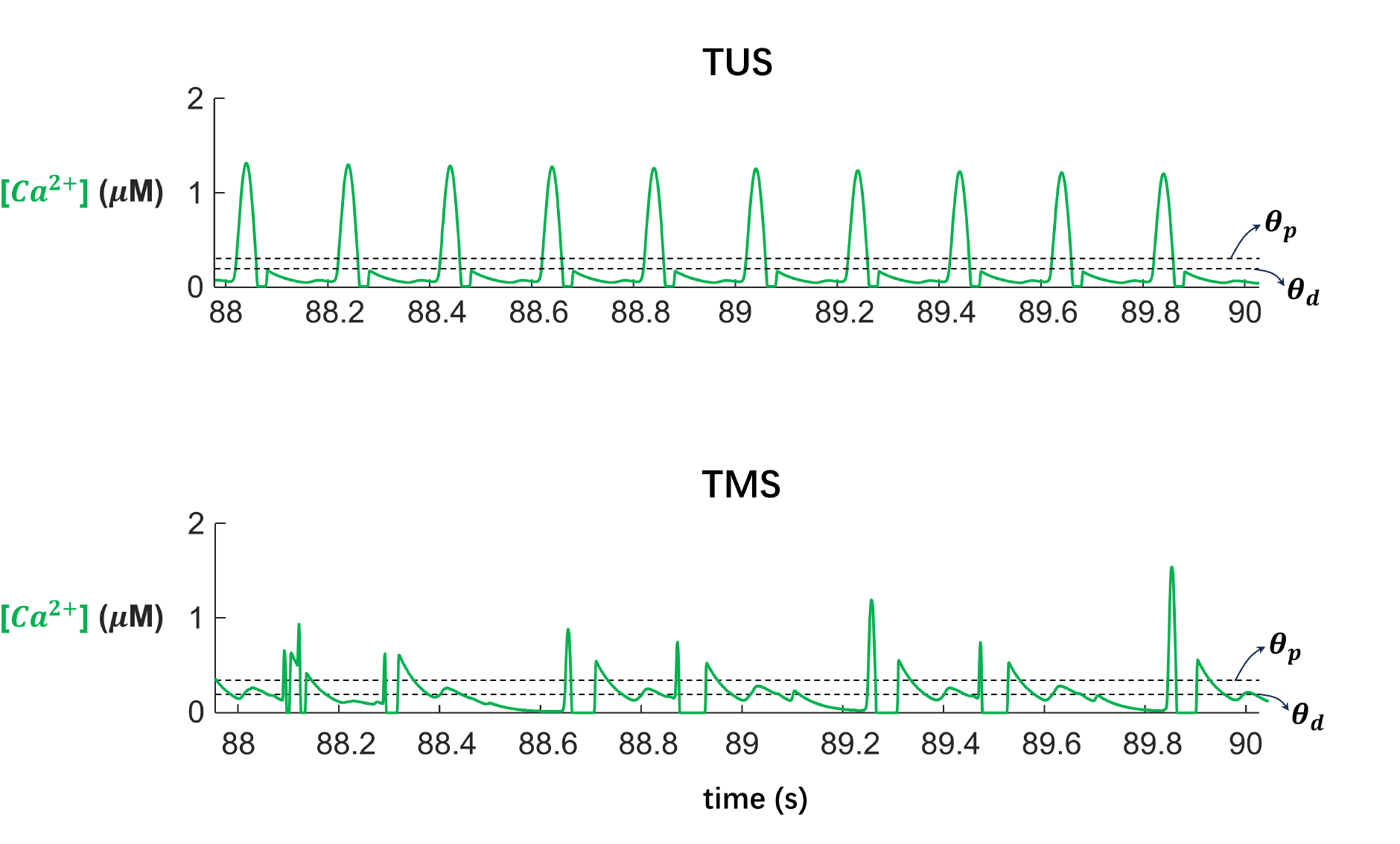
Calcium transients induced by TUS and TMS pulses. We compare the intracellular calcium concentration [*Ca*^2+^] transients induced by the TUS protocol cTB-TUS-40s and the TMS protocol cTB-TMS-40s. In both simulations, stimulation onset occurs at 50s; therefore, the figure displays the final two seconds of simulations. *θ*_*d*_ and *θ*_*p*_ denote the depression and potentiation thresholds, respectively, for the plasticity-induction Ω-function (**Eqs. 14-17, Fig. 3**).

**Figure S3:**
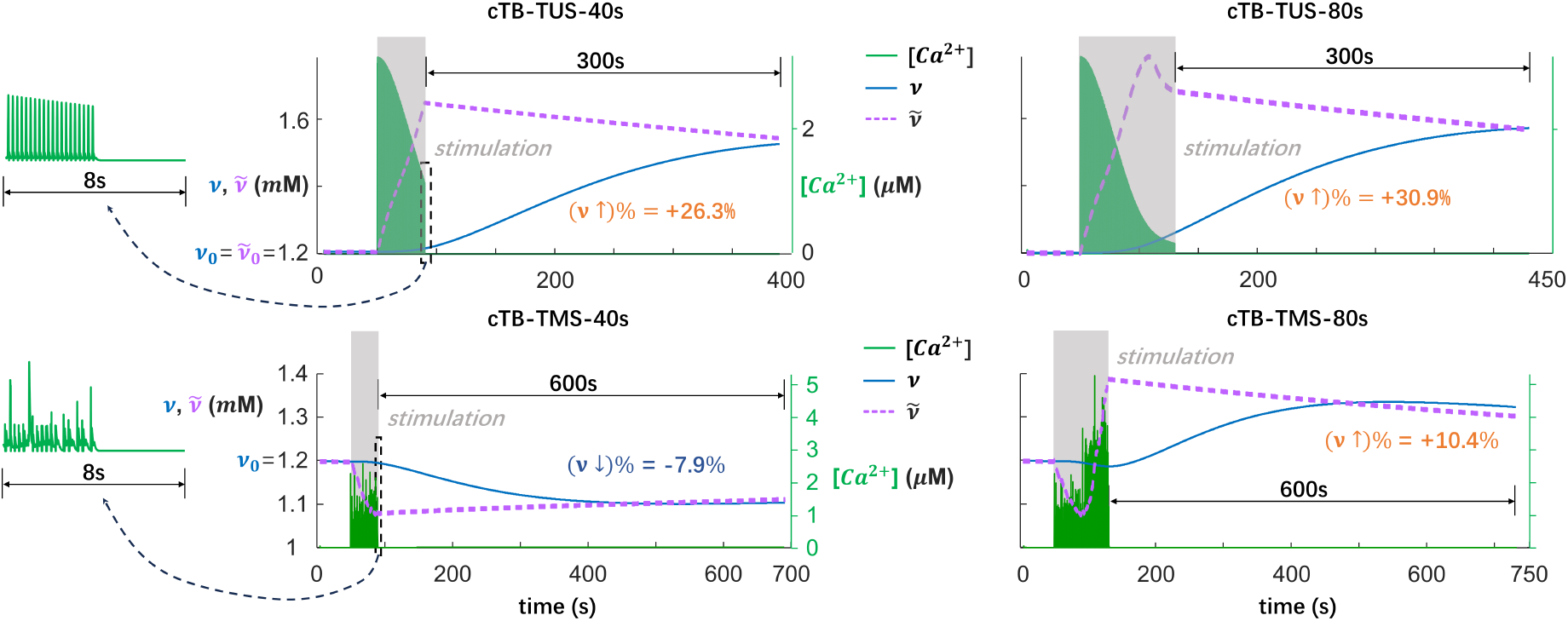
Complete time series of representative simulations in response to TUS and TMS. We present the complete simulation time series corresponding to the truncated traces shown in **Fig. 3**, where the displayed segments were selected to emphasize the calcium dynamics during stimulation.

